# Expanding the *MECP2* network using comparative genomics reveals potential therapeutic targets for Rett syndrome

**DOI:** 10.1101/2021.02.14.431162

**Authors:** Irene Unterman, Idit Bloch, Simona Cazacu, Gila Kazimirsky, Benjamin P. Berman, Bruria Ben-Zeev, Chaya Brodie, Yuval Tabach

## Abstract

Inactivating mutations in the Methyl-CpG Binding Protein 2 (MECP2) gene are the main cause of Rett syndrome (RTT). Despite extensive research into MECP2 function, no treatments for RTT are currently available. Here we use an evolutionary genomics approach to construct an unbiased MECP2 gene network, using 1,028 eukaryotic genomes to prioritize proteins with strong co-evolutionary signatures with MECP2. Focusing on proteins targeted by FDA approved drugs led to three promising candidates, two of which were previously linked to MECP2 function (IRAK, KEAP1) and one that was not (EPOR). We show that each of these compounds has the ability to rescue different phenotypes of MECP2 inactivation in cultured human neural cell types, and appear to act on Nuclear Factor Kappa B (NF-κB) signaling in inflammation. This study highlights the potential of comparative genomics to accelerate drug discovery, and yields potential new avenues for the treatment of RTT.

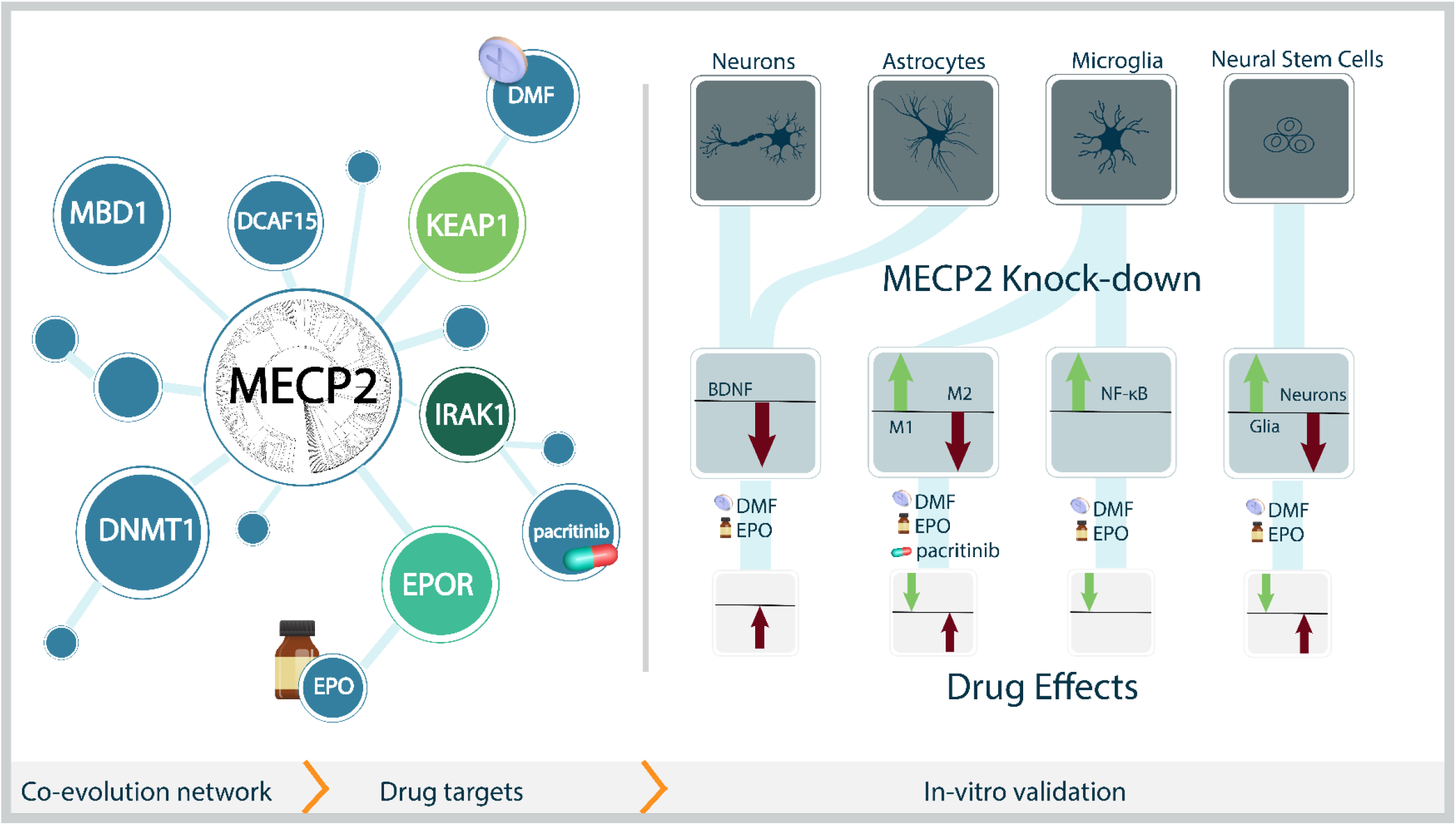

## Introduction

Rett syndrome (RTT [MIM: #312750]) is a rare genetic disorder caused by mutations in the methyl-CpG binding protein 2 (*MECP2*) gene *(1)*. It is an X-linked dominant disorder, almost exclusively affecting females. RTT is characterized by normal early development followed by regression, loss of purposeful hand movements, and intellectual disability *(2)*. Characteristics of the disease such as age of onset, range of symptoms, and their severity vary between patients *(3)*. The MECP2 protein has a role in transcriptional regulation but its exact mechanism of action, co-factors and downstream targets, are not fully characterized *(4)*.

The MECP2 protein, as a transcriptional regulator, has been linked to both gene silencing and activation *(5)*. It binds methylated CG dinucleotides through its methyl-binding domain and has a transcriptional repression domain that mediates binding to co-repressor proteins *(6)*. These two domains are hotspots for missense, nonsense and frameshift mutations for RTT disease *(7, 8)*. Transcriptomic analyses of RTT patient post-mortem brain tissues, blood samples, cell lines and murine models *(9)* revealed hundreds of affected genes but with low overlap between the studies. Thus the understanding of MECP2 mechanism of action, co-factors and downstream targets, is currently very limited *(4)*.

Therapies aiming to restore *MECP2* loss of function are in early development stages and are not approved for patients. These include attempts at read-through inducing drugs to overcome nonsense mutations *(10, 11)*, X reactivating therapies to allow expression of silenced endogenous *MECP2 (12)*, and gene therapy treatments to introduce wild-type *MECP2* copies intro the brain *(13, 14)*. Furthermore, current gene therapy methods show limited transduction efficiency and are unable to modify all cells in the target tissue *(15)*. Targeting of factors downstream of MECP2 is a promising avenue which could allow for partial amelioration of Rett symptoms, without the severe neurological phenotypes associates with MECP2 overexpression *(16, 17)*.

One major downstream target of MECP2 is brain-derived neurotrophic factor (BDNF) *(18)*, a key modulator of neuronal development and function. BDNF levels are reduced in symptomatic MECP2 KO mice *(19)* and experimental interventions that increase BDNF levels improve RTT-like phenotypes *(20)*. The IGF-1 protein potentiates BDNF activity, and treatments with recombinant human IGF-1 and its analogs have led to improvement in several clinical measurements *(21, 22)*, with a trial of an IGF-1 synthetic analog currently in Phase 3 (NCT04181723). Additional trials include the administration of triheptanoin supplementation which is being investigated in a Phase 2 clinical trial (NCT02696044) to target mitochondrial dysfunction in RTT*(23)*. Other interventions, such as ketamine (NCT03633058), targeting N-methyl-D-aspartate receptor (NMDAR) dysfunction, and Anavex 2-73 (NCT03758924), aimed at restoring mitochondrial dysfunction *(24)*, are currently being studied. The lack of understanding of pathways downstream of MECP2 emphasizes the need for accurate and comprehensive mapping of the MECP2 network with a focus on targetable proteins that could be used for interventions *(25)*. Here, we combine an innovative gene network approach (phylogenetic profiling) with drug targeting databases to construct such a map.

A well validated approach to the unbiased prediction of gene function and identification of functional gene networks is phylogenetic profiling (PP). The PP of a gene describes the pattern of conservation (i.e. presence or absence), of its orthologs in a set of genomes *(26)*. It relies on the assumption that if two or more genes share a similar phylogenetic profile then they may be functionally related. In our recent work, we developed an improved version of PP that normalizes signals across hundreds of eukaryotic genomes to identify important new members of cellular and disease pathways *(27–31)*. As protein function and evolution are very complex *(32)*, co-evolution within a local lineage (i.e. mammals, vertebrates, fungi, etc.) can provide complementary evidence. This is especially informative for genes with multiple functions that show complex or recent evolution *(27)*.

There are thousands of known bioactive compounds collected in databases such as Drug-Gene interaction database (DGIdb) *(33)* and Open Targets *(34)*, that contain information about validated direct or indirect effect on specific proteins or pathways. Network-based drug discovery methods are emerging as important tools to identify novel drug targets and predict the drug’s mode of action and potential side-effects *(35)*, and have been associated with higher success rates in clinical trials *(36, 37)*. Here, we perform phylogenetic profiling (PP) to construct a network of *MECP2* interacting genes that encode proteins targetable by known compounds. From this network, we selected three proteins that showed robust co-evolution with MECP2 across multiple evolutionary scales, had related roles in inflammation, and could be targeted by pharmacologically-validated compounds. These three compounds - pacritinib, EPO and DMF - were then functionally validated in *MECP2* Knock-Down (KD) human neural cells, including microglia, astrocytes and neural stem cells. The ability of all of these drugs to reverse some aspects of the MECP2-KD phenotypes demonstrates how comparative genome analysis can be used to understand and target this disease network.

## Results

### Phylogenetic profile analysis of MECP2 in eukaryotes and mammals identifies MECP2 co-evolved genes

To trace the evolutionary history of *MECP2*, we used BLASTP *(38)* to search the MECP2 protein against the proteomes of 1,028 eukaryotic species (see methods). We repeated the process for each of 20,192 human transcripts (longest transcript per gene) to generate a profile of conservation for each human gene. The scores were normalized according to sequence length and the evolutionary distance between humans and the queried species *(29)*. This Normalized Phylogenetic Profile (NPP) describes how conserved a transcript is in each species across the tree of life, compared to its expected conservation. It has been shown that genes with similar NPPs (i.e. genes that are either conserved or lost as a group) are functionally related, often belonging to the same pathway *(27–29, 31)*. We have shown that both across-clade and within-clade measures of co-evolution can be informative *(27, 32, 39)*, and thus we computed NPPs both across all eukaryotes (Fig. 1A and Fig. S1A) and within mammals (Fig. 1B and Fig. S1B). It is clear from the MECP2 NPP (Fig. 1A) that it is well conserved across mammals. Additionally, its expression patterns in mice are similar to human and it is essential for neuronal development in mice *(40)*.

**Figure 1.**
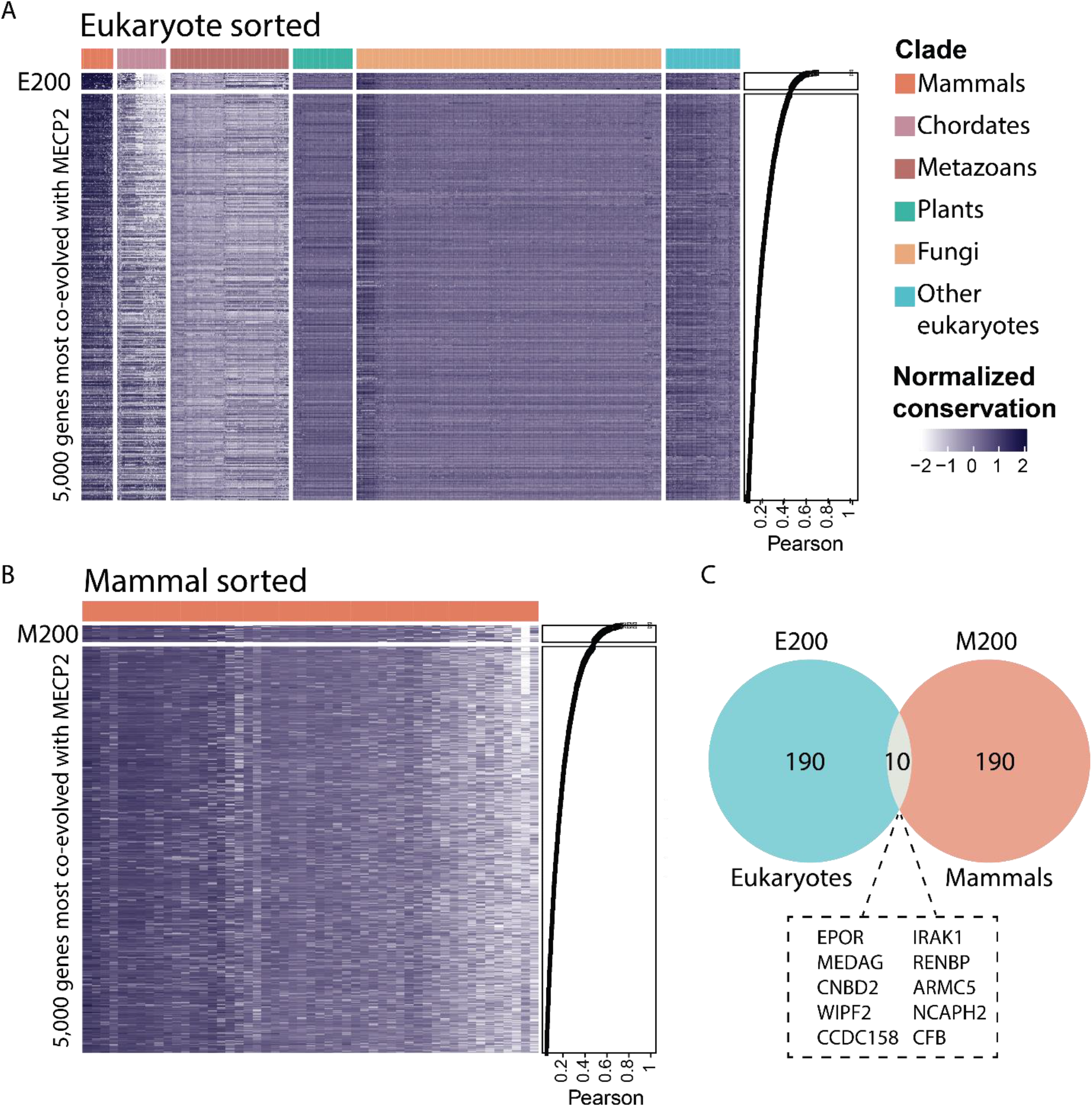
The phylogenetic profile of MECP2. A) The Normalized Phylogenetic Profile (NPP) of *MECP2* in eukaryotes along with the top 5,000 coevolved proteins. Each row represents one human protein, ordered by Pearson correlation to *MECP2*, with the top 200 proteins labeled as E200. Darker color indicates higher normalized conservation of the protein in each species. Organisms are grouped by phylogenetic clade, and clustered within each clade. B) The NPP of *MECP2* in 51 mammals along with the 5,000 top coevolved proteins, ordered by Pearson correlation to MECP within mammals. The top 200 proteins are labeled as M200. C) Gene overlap of the top 200 most co-evolved genes in eukaryotes and mammals, listing the 10 gene names listed in both.

We first used NPP scores to sort all human genes by their phylogenetic similarity to MECP2 across eukaryotes (Fig. 1A and Fig. S1A). We selected the top 200 genes (the “E200” set), which showed high correlation (R>0.47) with *MECP2*. Similarly, we used NPP scores within the mammalian clade (Fig. 1B and Fig. S1B) to select the top 200 mammal-centric genes (the “M200” set), which also showed high *MECP2* correlations (R>0.49). The E200 and M200 lists had 10 genes in common (Fig. 1C), representing a highly significant degree of overlap (p<3.2E-4 by the hypergeometric test). We considered the set of 10 overlapping genes, as well as the union set of 390 genes, as candidates for further exploration.

### MECP2 druggable protein network identifies several drug targets located in evolutionarily conserved chromosomal clusters

We chose to focus on the subset of *MECP2* co-evolved genes that could be targeted by readily available compounds. This approach not only allows for straightforward functional testing in both *in vitro* and animal models of *MECP2* inactivation, but moves us closer to the ultimate goal of identifying a candidate therapeutic for RTT. Using DGIdb *(33)* and Open Targets *(34)*, we searched specific drug-gene interaction types (e.g. inhibitory, activating) to identify direct effects on the 390 candidate genes. Open Targets identified such compounds for 11 proteins from our list, and DGIdb identified an additional 22, resulting in 33 druggable proteins in the MECP2 phylogenetic network.

Interestingly, two of these 33 proteins (IRAK1 and EPOR) were linked to MECP2 in both the eukaryotic E200 and the mammalian M200 lists, and both are targeted by compounds with clinical efficacy and acceptable safety profiles. Since co-evolved proteins have higher connectivity and tend to segregate within established functional networks *(28)*, we investigated these 33 druggable candidates using a STRING *(41)* interaction graph (Fig. 2A). IRAK1 was one of the six proteins functionally linked to MECP2, whereas EPOR had no known functional links. Because IRAK1 is known to be located on an evolutionarily conserved chromosomal domain with MECP2 (Fig. 2B and Fig. S2), we searched for other chromosomal domains enriched for MECP2 co-evolved genes. We identified the chr19p13.2 region as the most strongly enriched (p<1E-5) (Fig. 2C). This 5.6Mb domain contained 5 of the 33 druggable MECP2 coevolved genes - EPOR, DNMT1, ICAM1, ICAM3 and KEAP1 (Fig. 2D and Fig. S3). The MECP2 karyotype band Xq28 and chr19p13.2 were the only two bands in the genome that had more than one gene (Fig. 2C). Topologically Associated Domains (TADs) can mark functionally co-regulated gene clusters *(42)*, and the entire chr19p13.2 gene cluster is collocated within a TAD from high-resolution Hi-C mapping *(43)*. (Fig. 2E). Additionally this region has been shown to be bound by MECP2 *(44)*.

**Figure 2.**
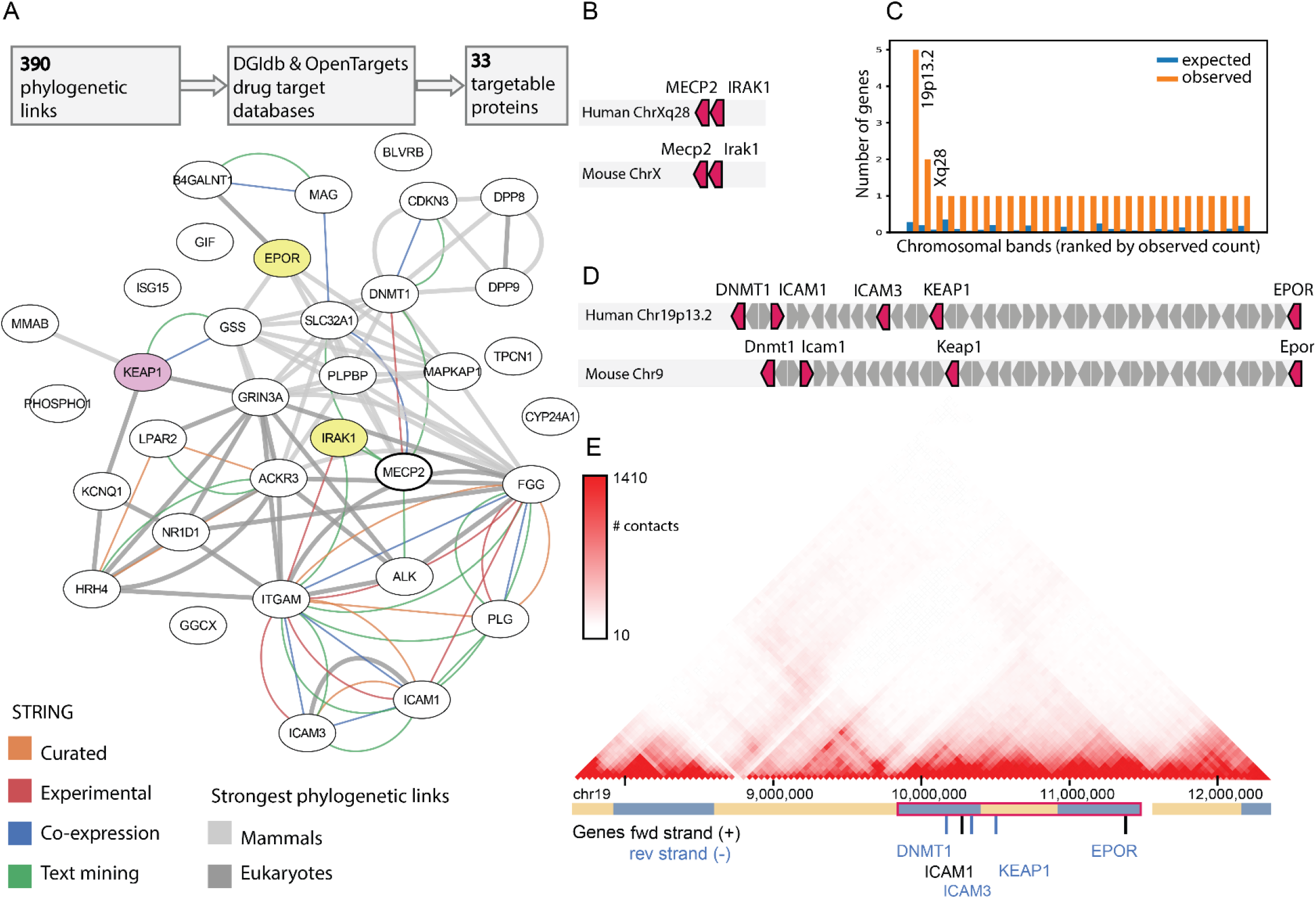
The *MECP2* druggable genes network. A) The 390 genes co-evolved with *MECP2* in either eukaryotes or mammals contain 33 genes that are targets of known drugs/compounds. These genes are shown in a STRING interaction graph, with known gene-gene interactions shown as colored edges, and those with very strong co-evolution (Pearson correlation > 0.7) shown as gray edges. The two proteins co-evolved in both eukaryotes and mammals (EPOR and IRAK1) are colored yellow. B) Genomic location of MECP2 interacting genes along chromosome X in humans and mice. C) Karyotype band locations of MECP2 and the 33 drug targets. D) Genomic location of MECP2 interacting genes along chr19p13.2 in humans and chromosome 9 in mice. E) Intra-chromosomal Hi-C contact heatmap for the chr19p13.2 locus in GM12878 cells, adapted from the 3D Genome Browser *(45)*. Genes in the MECP2 network are shown, with other genes hidden for clarity. TADs called by 3D Genome Browser shown in blue/yellow track, with super-TAD indicated containing MECP2 genes indicated.

For this list of six MECP2 co-evolved proteins located in evolutionarily conserved chromosomal clusters, we next investigated which could be targeted by an existing drug with known safety profile and efficacy in other contexts. NF-κB signaling has previously been implicated in RTT, with heterozygous deletion of *NFKB1* dramatically increasing lifespan in *MECP2* null mice *(46)*. IRAK1 was identified as a key intermediary in this process, with MECP2 directly binding the IRAK1 promoter to regulate its expression in MECP2-null mouse brains *(46)*. Pacritinib is a JAK2/FLT3 inhibitor with IRAK1 inhibiting capabilities *(47)*, presently under clinical investigation for myelofibrosis and glioblastoma. As it is currently the only clinical stage IRAK1 inhibitor with known clinical efficacy and acceptable safety even after prolonged administration *(48)*, we chose this compound for our functional studies.

EPOR forms a heterodimer with β common receptor (βCR), also known as the Tissue-Protective Receptor (TPR) *(49)*. The endogenous hormone EPO binds the TPR activating signaling cascades with roles in neuroprotection *(50)*. EPO has a well understood pharmacological profile, and several variants of erythropoietin lacking hematopoietic effects have been developed and shown to preserve the neuroprotective effects of EPO administration *(51, 52)*.

KEAP1 is the key repressor of the transcription factor nuclear factor erythroid 2-related factor 2 (NFE2L2), which is involved in response to antioxidants through its transcriptional regulation of inflammatory pathways. Dimethyl fumarate (DMF) is an approved treatment for both multiple sclerosis and psoriasis *(53, 54)*. Its administration decreases KEAP1 levels, increases NFE2L2 levels and causes dissociation and nuclear transport of NFE2L2. DMF treatment has proven to be effective and safe in conferring neuroprotection *(55)*.

These three candidate proteins targeted by promising drug products (IRAK1, EPOR, and KEAP1) all suggested possible roles mediating inflammation within the MECP2 network. Therefore, we tested these three drug candidates (Pacritinib, EPO, and DMF) in neural cell models and inflammatory phenotypes.

### Human neural cell cultures exhibit Rett-like phenotypes when MECP2 is silenced

To analyze the effects of the three selected compounds on a RTT model system, we employed human immortalized primary neural cells silenced for MECP2. We chose to use a human *in-vitro* model system for preliminary validation due to the significant differences in the phenotypes and functions of mouse and human neural cells *(56)*. Indeed, recent studies reported the use of iPSCs and immortalized glial cells as reliable human models for analyzing specific therapeutic targets for various diseases including autism *(57)*, neuroinflammation *(58, 59)* and for analyzing the crosstalk of glioma with glial cells *(60–62)*.

Since all three selected compounds were associated with changes in neuroinflammation, we first focused on microglia and astrocytes. Loss or impaired function of MECP2 has been reported to affect the functions of glial cells and these changes have been implicated in the pathogenesis of RTT *(63, 64)*. We first characterized the effects of MECP2 silencing on the phenotypes of microglia. For these experiments, we transduced the cells with lentivirus vectors expressing control or MECP2 shRNAs. The expression of MECP2 was silenced by over 80% in these cells (Fig. S5). MECP2 silencing induced a relative increase in the expression of M1 markers (IL1, and CD86) compared to M2 markers (CD206 and IL-13) (Fig. 3A), suggesting an increase in neuroinflammation. We also found that MECP2 silencing decreased the phagocytosis of the microglia cells (Fig. 3B).

**Figure 3.**
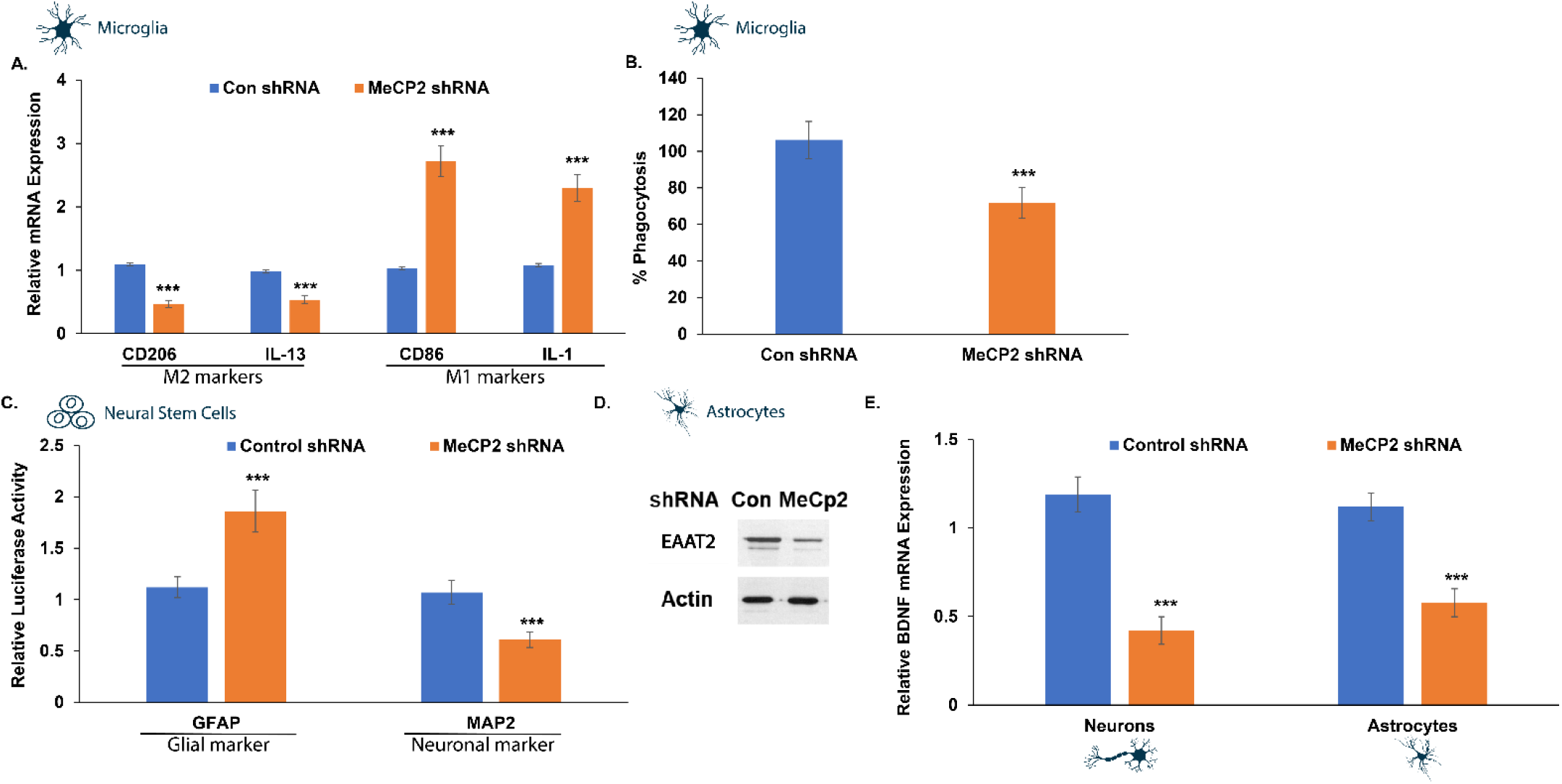
Effects of MECP2 knockdown on neural cell phenotypes. Human microglia (A,B), NSCs (C) astrocytes (D,E) and neurons (E), were silenced for M ECP2 using lentivirus vector. The relative expression of M1 and M2 markers was analyzed in microglia cells using RT -PCR (A) and degree of phagocytosis using the pHrodo^™^ assay (B). NSCs were transduced with lentivirus vectors expressing the differentiation reporters GFAP and MAP2 and were differentiated as described in the methods. 10 days later, luciferase activity was determined (C). Astrocytes silenced for MECP2 were analyzed for the expression of EAAT2 using Western blot analysis (D) and the expression of BDNF mRNA in both MECP2 silenced astrocytes and neurons was determined using RT -PCR. The results are the means ± SD of a representative experiment of three separate tests analyzed in quadruplet. *** P<0.001

We next analyzed the effect of MECP2 silencing on human astrocyte differentiation and functions. For these experiments, neural stem cells (NSCs) were transduced with lentivirus vectors expressing the differentiation reporters, GFP/fLuc GFAP or the mCherry/Luc MAP2. The cells were plated on laminin coated plates and luciferase activity was determined 10 days later. In accordance with previous reports, we found that MECP2 silenced NSCs displayed preferential expression of the glial differentiation marker (GFAP) with a concomitant decrease in expression of the neuronal differentiation marker (MAP2) (Fig. 3C). The MECP2 silenced astrocytes expressed lower levels of Excitatory Amino Acid Transporter 2 (EAAT2), indicating impaired function of these cells, as was suggested for mouse astrocytes. (Fig. 3D).

We finally examined the effects of MECP2 silencing on the expression of BDNF in human neurons and astrocytes and found that, as reported for mouse cells, MECP2 silencing decreased the expression of BDNF mRNA in both astrocyte and neuronal cultures (Fig. 3E). Altogether, these finding indicate that the immortalized human cells employed in this study represent a reliable model for studying MECP2-related pathways and potential treatments.

### DMF, EPO and pacritinib differentially abrogate the effects of MECP2 silencing on microglia polarization

We then analyzed the effects of the three selected compounds, DMF, EPO and pacritinib, on MECP2 silenced cells. We first demonstrated that in the concentrations range we used, none of these compounds exerted a toxic effect on neither microglia cells, astrocytes, primary neuronal cells nor NSCs (Fig. S4).

Control and MECP2 silenced microglia were treated with the DMF, EPO and pacritinib and their effects on the relative expression M1 (IL1, TNFα and CD86) and M2 (CD206 and IL-13) markers were determined using RT-PCR. The three compounds abrogated the M1 shift induced by MECP2 silencing, albeit to a different degree. EPO exerted the most significant effect and increased the expression of both M2 markers, CD206 and that of IL-13, while decreasing the expression of the M1 markers, CD86, IL-1 and TNF-α (Fig. 4A). In contrast, both DMF and pacritinib exerted only partial effects on the MECP2 silenced cells and abrogated the inhibitory effects of MECP2 silencing on the expression of CD206, CD86. Whereas, they did not exert a significant effect on the decreased IL-13 expression (Fig. 4B, Fig. 4C). Similarly, both DMF and pacritinib decreased the expression of the M1 marker TNF-α (Fig. 4B) but not that of IL-1 (Fig 4B, Fig. 4C). We also analyzed the effects of the different compounds on the phagocytosis of MECP2 silenced microglia cells. Treatment of MECP2 silenced cells with DMF exerted the strongest increase in the phagocytosis of the silenced cells, whereas EPO and pacritinib did not exert a significant effect (Fig. 4D).

**Figure 4.**
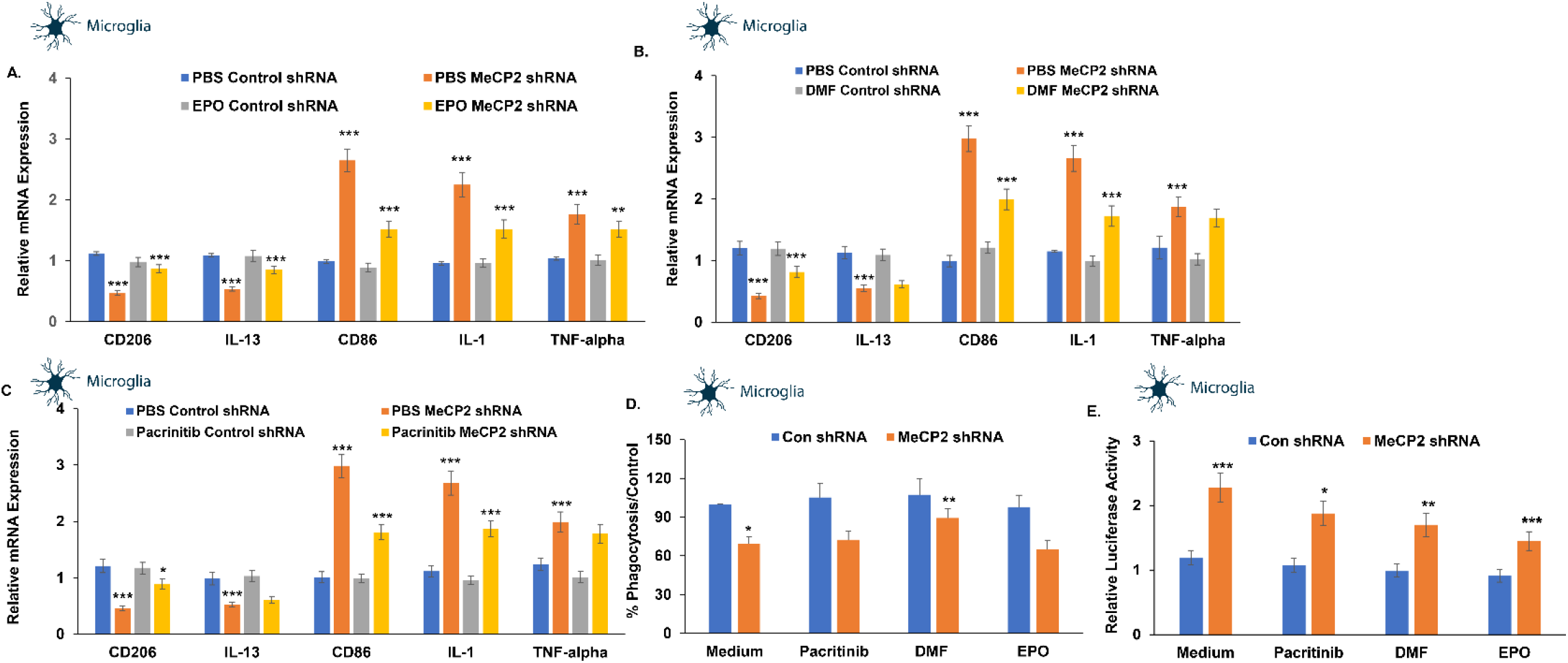
Effects of tested compounds on the polarization of microglial cells. Human microglia cells were silenced for MECP2 using lentivirus vectors expressing MECP2 shRNA. Control cells were transduced with lentivirus vectors expressing a control shRNA. After five days, the expression of MECP2 was determined by Western blot analysis (Fig. S5). Control and silenced cells were treated with EPO 10 ng/ml (A), DMF 10 uM (B) and pacritinib 10 uM (C), and the expression of M1 and M2-associated markers were determined after 72 hours using RT-PCR. MECP2 silenced microglia cells treated with EPO, DMF or pacritinib were also analyzed for phagocytosis using the pHrodo^™^ assay (D). MECP2-silenced microglia cells were transduced with lentivirus vectors expressing the NF-kB reporter followed by treatment with EPO, DMF and pacritinib for 24 hours. Luciferase activity was determined (E). The results demonstrate the means ± SD of a representative experiment of three separate tests analyzed in quadruplet. *** P<0.001 PBS Con shRNA vs PBS MECP2 shRNA; MECP2 silenced cells, treated cells vs. untreated for EPO: CD86 and IL- 1 expression; DMF and pacritinib: CD86 expression and NF-kB activation of EPO-treated cells. **P<0.01 DMF: IL- 1 and phagocytosis, pacritinib: CD206 and IL-1 and NF-kB activation of DMF-treated cells.* P<0.05, EPO: IL-13 and TNF-α and NF-kB activation of pacritinib-treated cells

### EPO and DMF inhibit NF-κB activation in MECP2 silenced microglia cells

Knockdown of MECP2 has been reported to lead to increased NF-κB activity in neuronal and myeloid cells *(65)* and this deregulated activity has been associated with neuroinflammation and decreased dendritic arborization and spine density in a mouse model of RTT *(46)*. We therefore examined the ability of our three candidate compounds to abrogate the increased NF-κB activity in MECP2 KD microglia cells. MECP2-silenced microglia cells were transfected with lentivirus vectors expressing the NF-κB luciferase reporter and a constitutively active Renilla luciferase construct (as a transfection control). The cells were treated with EPO, DMF and pacritinib for 24 hours and luciferase activity was determined thereafter. A significant increase in NF-κB-dependent luciferase activation was observed in the MECP2 silenced cells (Fig. 4E). This increase was abrogated in cells treated with EPO and DMF and to a lesser degree by pacritinib, indicating that EPO and DMF were able to downregulate the increased NF-κB activation in MECP2 KD cells.

### EPO affects the astrocytic differentiation and function of MECP2-silenced NSCs and astrocytes

Various studies support a non-cell-autonomous effect of astrocytes on neuronal cell functions and contribution of astrocytes to various aspects of Rett syndrome pathogenesis via regulation of glutamate levels, homeostasis and neuroinflammation *(63, 66)*. As presented in Figure 3C, MECP2 silencing increased astrocytic differentiation of NSCs at the expense of neuronal differentiation. Treatment of the silenced NSCs with pacritinib or DMF did not have significant effects on the differentiation of these cells. In contrast, EPO abrogated the impaired NSC differentiation induced by MECP2 silencing and increased neuronal differentiation (Fig. 5A). As demonstrated in Figure 3D, MECP2 silencing decreased the expression of the glutamate transporter EAAT2 in human astrocytes (Fig. 5B). Treatment of the cells with EPO inhibited the decreased EAAT2 expression as demonstrated by RT-PCR, whereas pacritinib and DMF did not have a significant effect (Fig. 5B).

**Figure 5.**
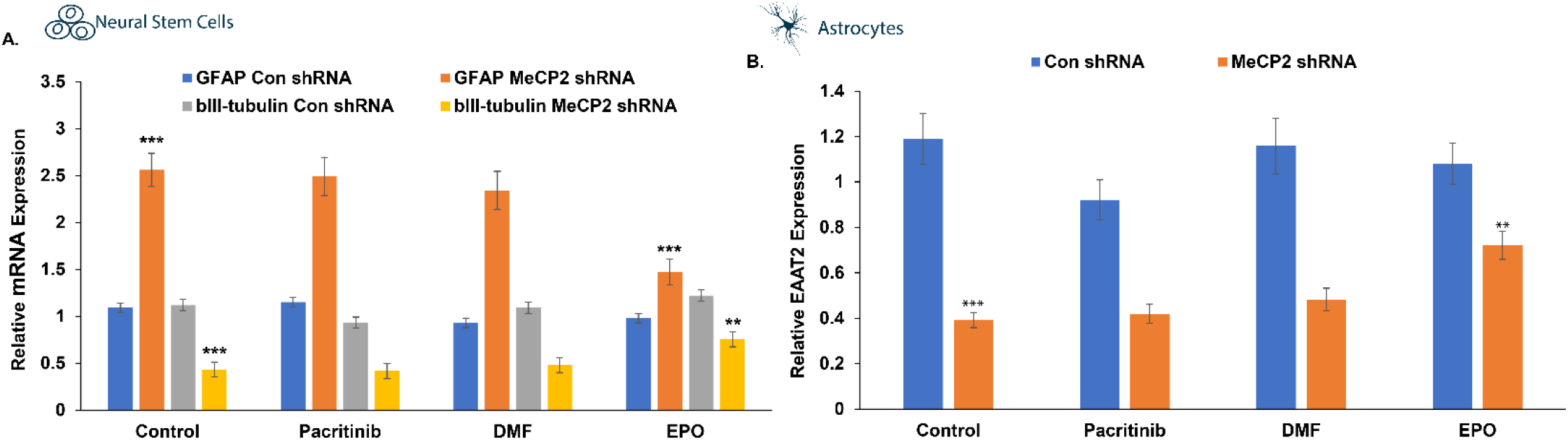
Effects of tested compounds on the differentiation of NSCs and EAAT2 expression in astrocytes. Human NSCs (A) and astrocytes (B) were silenced for MECP2 using lentivirus vectors expressing MECP2 shRNA. Control cells were transduced with lentivirus vectors expressing a control shRNA. After five days, the expression of MECP2 was determined by Western blot analysis (Fig. 5S). A) Control and silenced NSCs were allowed to differentiate for 10 days and the expression of GAFP and βIII tubulin were determined using RT-PCR. B) Silenced astrocytes treated with the different compounds or with medium were analyzed for the expression of EAAT2 using RT-PCR. The results are the means ± SD of a representative experiment of three separate tests analyzed in quadruplet.

### DMF and EPO upregulate BDNF expression in MECP2-silenced cells

BDNF plays important roles in neuronal growth and development and represents a well-recognized transcriptional target of MECP2 *(18)*. Silencing of *MECP2* decreased BDNF expression in both neurons and astrocytes. DMF and EPO induced an increase in BDNF mRNA in the silenced neurons (Fig. 6A) and in BDNF secretion in the silenced astrocytes (Fig. 6B), whereas pacritinib did not exert a significant effect.

**Figure 6.**
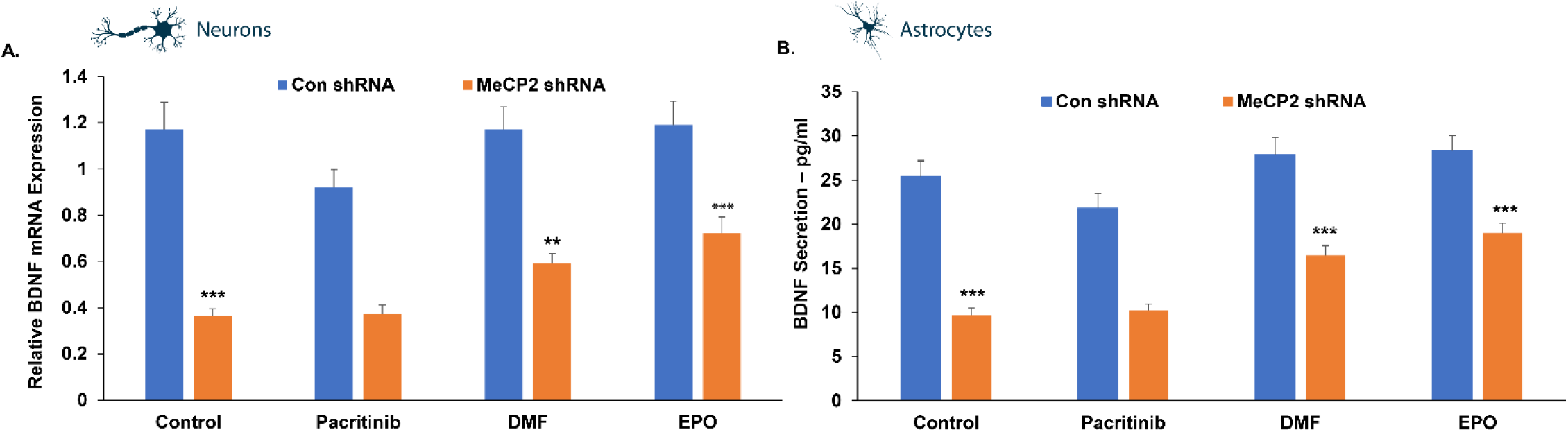
BDNF expression in neural cells. MECP2 expression was silenced in human neurons (A) and astrocytes (B) and BDNF expression was analyzed in cells treated with a medium or the specific tested compounds using both RT-PCR (A) and ELISA (B). The results are the means ± SD. ***P<0.001

## Discussion

Rett syndrome is a neurodevelopmental disease without evidence of degeneration and thus rescuing MECP2 downstream activity might improve disease pathophysiology even in adulthood *(67)*. In murine models, reintroduction of *MECP2* to adult animals led to vast phenotypic improvements *(13)*. In parallel with approaches aimed at increasing MECP2 levels directly *(10, 11, 68)*, accurate mapping of the MECP2 gene network is important to provide additional avenues for intervention, and avoid the toxicity associated with overexpression of MECP2 *(17)*. Here, we have presented a framework for identifying potential drug-gene interactions based on comparative genomics and validate these interactions in human immortalized primary neural cells. We analyzed gene conservation across mammals and hundreds of eukaryotes to infer functional links and generated a network of co-evolved genes, identifying known targets such as IRAK1 as well as completely new candidates such as EPOR. We then mined this network to identify actionable drug targets, prioritizing a list of associated compounds using a combination of strong co-evolutionary evidence of the target genes and a proven safety profile of the pharmaceutical compounds (Pacritinib, DMF, and EPO).

We reproduced a number of *in-vivo* RTT phenotypes in human neural cell cultures with decreased MECP2 expression. When treated with the three selected compounds, these phenotypes could all be reversed to different degrees in a cell-dependent manner. While all the compounds restored the polarization of microglia, the decreased phagocytosis of these cells was abrogated only by DMF treatment. In these cells, DMF, EPO and pacritinib differentially inhibited the hyperactivity of NF-κB signaling. DMF and EPO abrogated the inhibitory effect of *MECP2* silencing on BDNF expression and secretion, whereas pacritinib did not have a significant effect. EPO also partially reversed the impaired NSC differentiation and astrocytic EAAT2 expression in silenced NSCs and astrocytes, respectively.

The activity of these compounds on microglia polarization and NF-κB activity is particularly interesting. Activation of microglia cells has been reported in *MECP2* KO mice *(69)* and has been associated with pathological process in RTT *(63)*. An MECP2 KO mouse model also implicated NF-κB signaling and IRAK1 as important mediators of the neuroinflammatory response in RTT *(46)*. While the mechanisms involved in the reversing effects of these three compounds remains to be determined, NF-κB signaling downstream of MECP2 is an attractive candidate (Fig. 7). EPOR and KEAP1 have also been implicated as NF-κB modulators involved in the inflammatory response. The endogenous hormone EPO binds the TPR to suppresses inflammatory cytokines in immune cells, a process which involves inhibition of the NF-κB pathway through GSK3β *(52)*. NFE2L2, usually bound by KEAP1, also has a direct cytoplasmic role in modulating NF-κB signaling through degradation of IκB and nuclear translocation of NF-κB *(70)*. The three compounds tested all inhibited NF-κB-dependent luciferase activation in microglia cells to a certain extent, validating their involvement in this context.

**Figure 7.**
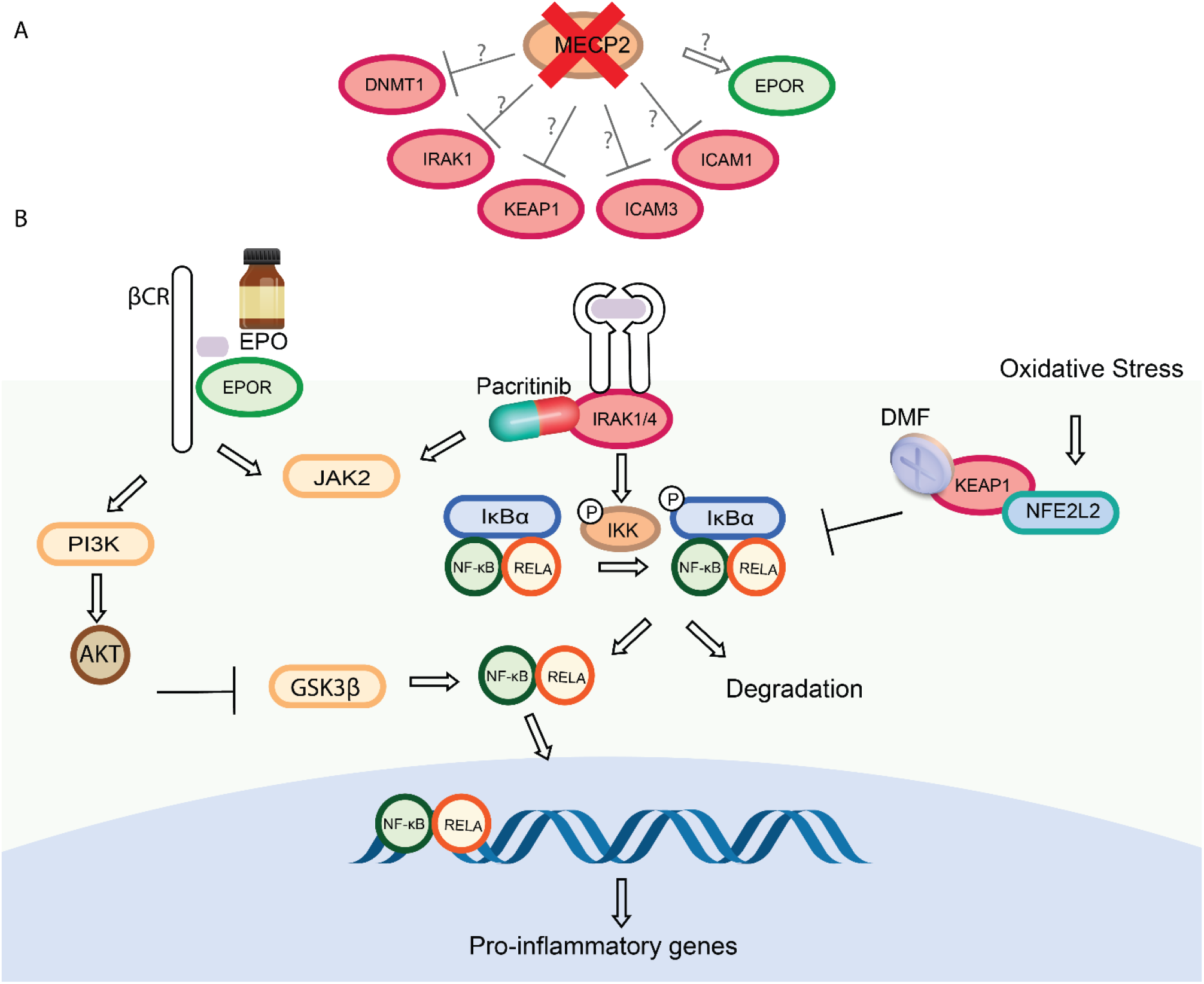
MECP2 network genes converge on NF-κB signaling. (A) We propose that reduced MECP2 levels could lead to increase in ICAM1, ICAM3, IRAK1, and KEAP1 levels, and a decrease of EPOR levels within the tissue-protective receptor. (B) IRAK-1 plays a role in NF-κB activation through IKK activation, which leads to IκB phosphorylation and subsequent degradation, NF-κB nuclear translocation and the expression of its pro-inflammatory target genes. KEAP1 binds NFE2L2 and prevents its nuclear localization. NFE2L2 target genes inhibits NF-κB signaling through a non-transcriptional mechanism involving degradation of IκBα. EPO binds EPOR and enhances the activation of Akt, resulting in inhibition of GSK-3β and inhibition of NF-κB nuclear transport.

In addition to the inflammatory effect, we found that *MECP2* silencing decreased the neuronal while increasing the astrocytic differentiation of NSCs. However, human astrocyte function was impaired in the MECP2 silenced cells and these cells expressed lower levels of the glutamate transporter EAAT2. The signaling pathways involved in these effects of EPO are not fully characterized, however, NF-κB has been recently reported to regulate EAAT2 expression in astrocytes *(71)*.

Finally, we found that both DMF and EPO rescued the decreased BDNF expression in MECP2-silenced microglia and astrocytes. Indeed, both NF-κB and NFE2L2 pathways have been reported to be associated with the regulation of BDNF expression *(72, 73)*. Thus, DMF can increase BDNF levels through an increase in NFE2L2, which binds the *BDNF* promotor and acts as a transcriptional activator *(73)*. EPO administration activates the STAT3 signaling pathway *(74)*, which can in turn elevate BDNF expression *(75)*. Recently, EPOR activation through rhEPO administration was reported to induce neuro-differentiation after hypoxia in mice *(76)*.

In summary, EPO, DMF and pacritinib act differentially on MECP2 silenced NSCs, microglia and astrocytes, providing a preliminary validation of our approach. The investigation into specific drug effects and mechanisms of actions are currently being conducted. In addition, *in-vivo* studies of EPO, DMF and pacritinib using MECP2 KO mice are underway to complement the *in-vitro* findings and assess whether any of the identified drugs can be further be explored for clinical use.

## Materials and Methods

### Mapping MECP2 conservation along 1,028 eukaryotic species and identifying correlated genes

We used the Normalized Phylogenetic Profile (NPP) *(28, 29)* of 1028 eukaryotic species to rank genes co-evolved with MECP2. Briefly, 20,192 human proteins (one representative protein sequence per gene) were downloaded from UniProt reference proteomes (June 2018 release) *(77)*. When different splice variants existed for a gene, the longest variant was used. The proteins were searched with BLASTP *(38)* against the proteomes of the 1028 eukaryotes. A bit-score of 20.4, corresponding to a BLAST e-value of 0.05, was set as a minimal similarity threshold. The top scoring protein in each organism was selected. The bit-scores were normalized to protein length and phylogenetic distance from humans. The output is a matrix P of size 20,242 × 1028 where each entry Pab is the best BLASTP bit score between a human protein sequence ‘a’ and the top result in organism ‘b’. Pearson correlation was calculated for each profile with the MECP2 profile. The top 200 genes in all eukaryotes were selected as the E200 group for further analysis. This procedure was repeated using only the 51 mammal species, and the top 200 genes in were selected as the M200 group for further analysis.

### Filtering genes with drug interactions and constructing protein network

Drug gene interactions were collected from DGIdb *(33)* and from Open Targets *(34)*. Drugs at any stage of development were included in the search as long as the interaction with the target gene had known directionality (e.g. inhibitory, activating). Drug-gene interactions were validated through a literature review and known gene-gene interactions were collected from STRING *(41)*. Network diagram was constructed using STRING and Cytoscape *(78)*.

### Synteny and co-localization analysis

Gene locations were retrieved from BioMart *(79)*, the NCBI genome data viewer (Fig. S2A, S3A), and Genomicus *(80)* (Fig 3A). Synteny in ancestral species was obtained from Genomicus *(80)*. Intra-chromosomal Hi-C contact heatmaps and TAD borders were collected from the 3D Genome Browser *(45)*, using GM12878 cell data from the high-resolution Hi-C dataset of Rao et. al *(43)*.

### Neural cell cultures

Immortalized human microglial cells and astrocytes were obtained from Applied Biological Material (Richmond, BC, Canada). Human NSCs (H9, hESC-derived) (ReNcell) were obtained from Invitrogen, Merck (Germany). Human neurons were obtained from ScienCell (Carlsbad, CA, USA). All cells employed in this study were tested for mycoplasma contamination (Mycoplasma PCR Detection Kit) and found negative.

### Transduction of neural cells

Lentivirus vectors (System Biosciences, Mountain View, CA, USA) expressing the MECP2 or control shRNAs and the reporters GFP/fLuc GFAP, mCherry/Luc MAP2 and NF-kB luciferase were packaged and used to transduce the cells according to the manufacturer’s protocol and as previously described *(81)*.

### NSC differentiation

Human NSCs were maintained as spheroids in an NSC maintenance medium containing fibroblast growth factor 2 (FGF-2) and an epidermal growth factor (EGF, 20 ng/ml) on laminin-coated flasks. For differentiation, the cells were maintained in NSC maintenance medium without FGF-2 and EGF and neuronal and glial differentiation were observed.

### Microglia polarization

Human microglia cells were silenced for MECP2 using lentivirus vectors expressing MECP2 or control shRNAs. After five days, the expression of MECP2 was determined by Western blot analysis and cells exhibiting a decrease of at least 80% were employed in further studies. Control and silenced cells were treated with the specific compounds and the expression of M1- and M2-associated markers were determined after 72 hr using RT-PCR. All experiments were done in triplicates and were repeated three times.

### Cytotoxicity assay

Cells were treated with different concentrations of DMF (1 10 and 50 mM), EPO (10, 30 and 50 ng/ml) and Pacritinib (1 and 10 mM). Following three days of treatment, the cells were analyzed for cell death using LDH assay. The results are presented as the means ± SD of six samples for each compound.

### Western blot analysis

Cell pellet preparation and Western blot analyses were performed as previously described *(82)*. Equal loading was verified using an anti-β-actin or tubulin antibodies as described *(81, 83)*.

### Real-time PCR

Total RNA was extracted using RNeasy midi kit according to the manufacturer’s instructions (Qiagen, Frederick, MD, USA). Reverse transcription reaction was carried out using 2-μg total RNA *(83)*. The primer sequences are described in the supplementary files (Table S1).

### Phagocytosis analysis

Human microglial cells were silenced for MECP2. Phagocytosis was determined using the pHrodo^™^ Green zymosan bioparticle assay (Invitrogen, Carlsbad, CA, USA) according to the manufacturer’s instructions. Briefly, microglia were incubated with a solution of pHrodo Green zymosan bioparticles in Live Cell Imaging Solution (0.5 mg/ml) for two hr. Phagocytosis was determined using a fluorescence plate reader at Ex/Em 509/533.

### Luciferase activity

The firefly luciferase activity of the NF-kB, GFAP and MAP 2 activities and the control Renilla luciferase activity were analyzed using the Dual-Luciferase® Reporter Assay System (Promega Corporation).

### Statistical analysis for cell assays

The results are presented as the mean values ± SD. Data were analyzed using analysis of variance or a Student’s t test with correction for data sets with unequal variances.

## Web resources

OMIM, http://www.omim.org/.

Genomicus, https://www.genomicus.biologie.ens.fr/

NCBI Genome Data Viewer, https://www.ncbi.nlm.nih.gov/genome/gdv/

STRING, https://string-db.org/

DGIDB, http://www.dgidb.org/

Open targets, https://www.opentargets.org/

Uniprot, http://www.uniprot.org/

Graphics include vectors designed by macrovector_official / Freepik, https://www.freepik.com/

## Acknowledgments

We thank Dr. Yaron Daniely and the Yissum team for being a major driving force for this project. We thank Dr. Liana Patt and the Integra Holdings team for their support. We thank Dr. Hae Kyung Lee and Susan Finniss for their technical support. We thank Dr. Amir Eden and Prof. Yinon Ben Neriah for their insight and comments.

## Author contributions

YT conceptualized and supervised the study. IU performed the phylogenetic analysis and drug selection. IB constructed the phylogenetic matrix used for analysis. CB, GK and SC established the human MECP2 KD cell system and performed all experiments. BPB contributed to result interpretation and drafting of manuscript. BBZ contributed to the study design and result interpretation. IU and CB wrote the original draft with contributions from the co-authors. All authors read and approved the final manuscript.

## Data and materials availability

The data that support the findings of this study are openly available in CladeOScope [at http://cladeoscope.cs.huji.ac.il/]. Code is available upon request.

## Supplementary Materials

**Figure S1.**
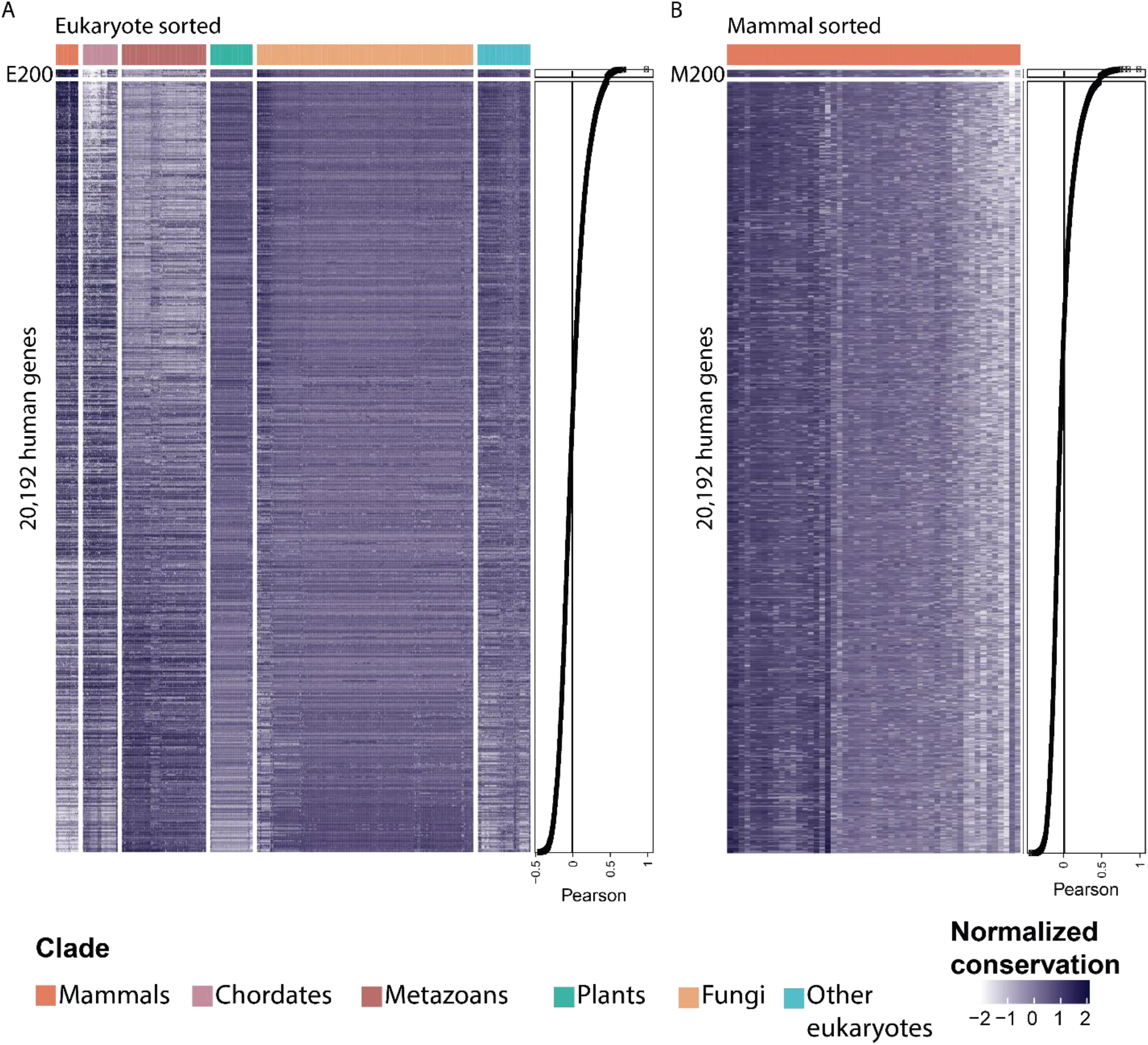
Scaled phylogenetic profiles of all human genes by similarity to MECP2. A) NPP profiles of human genes in 1028 eukaryote species, ordered by the Pearson correlation to the profile of MECP2. B) Phylogenetic profiles of human genes in 51 mammals, scaled to highlight the conservation patterns within the clade.

**Figure S2.**
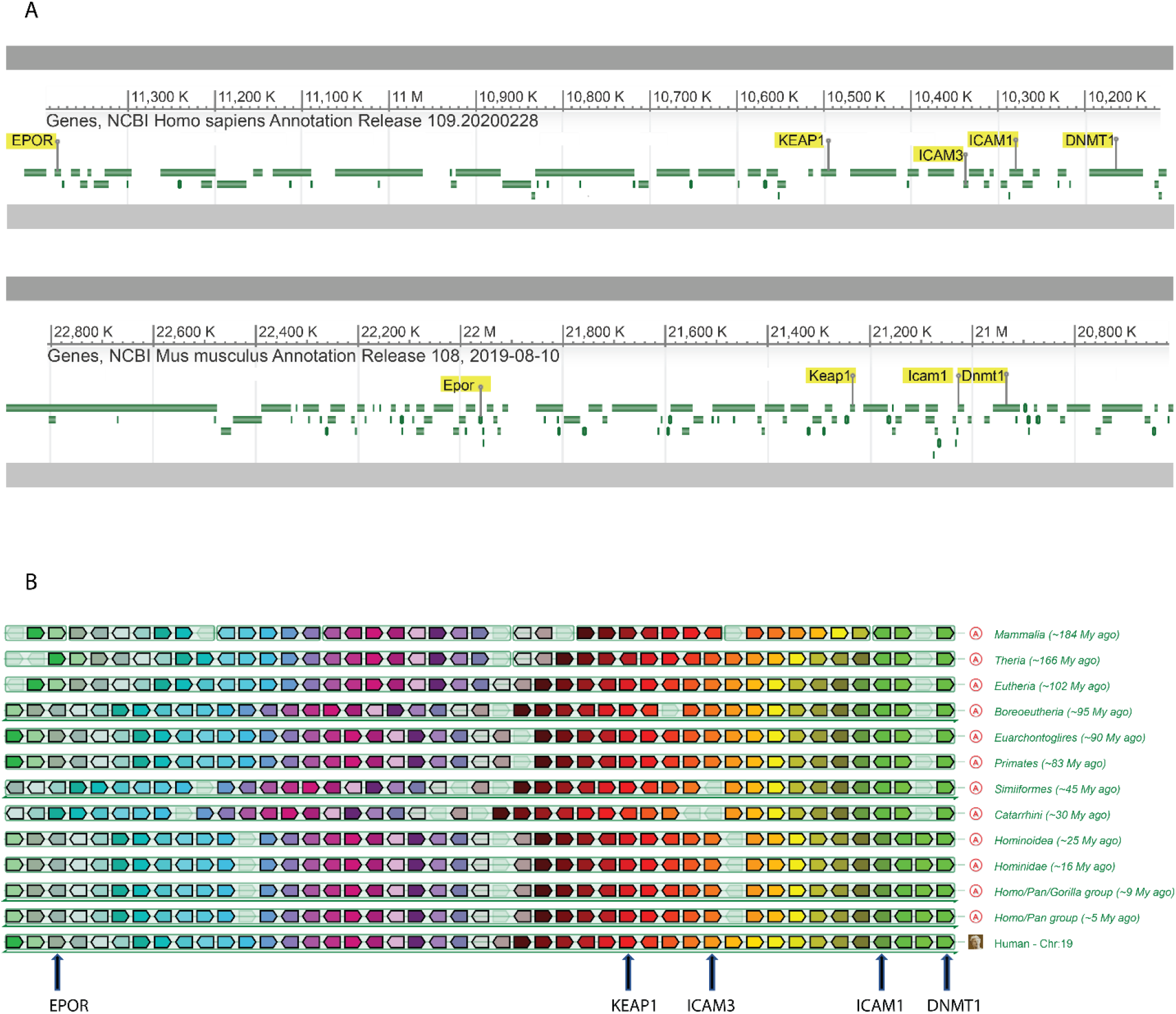
Synteny in the Chr19p13.2 locus. A) NCBI Genome Data Viewer view of the genomic locations of DNMT1, ICAM1, ICAM3, KEAP1 and EPOR (top panel, genes highlighted). The gene orthologs are shown in mus musculus chromosome 9 (bottom panel). B) Inference of the synteny in ancestral species from Genomicus. Each row represents the same genomic locus at an ancestral state. Each gene is represented by a different color, and selected genes are annotated below. EPOR, ICAM1, ICAM3, KEAP1 and DNMT1 maintain their relative positions as far as the mammalian ancestral genome.

**Figure S3.**
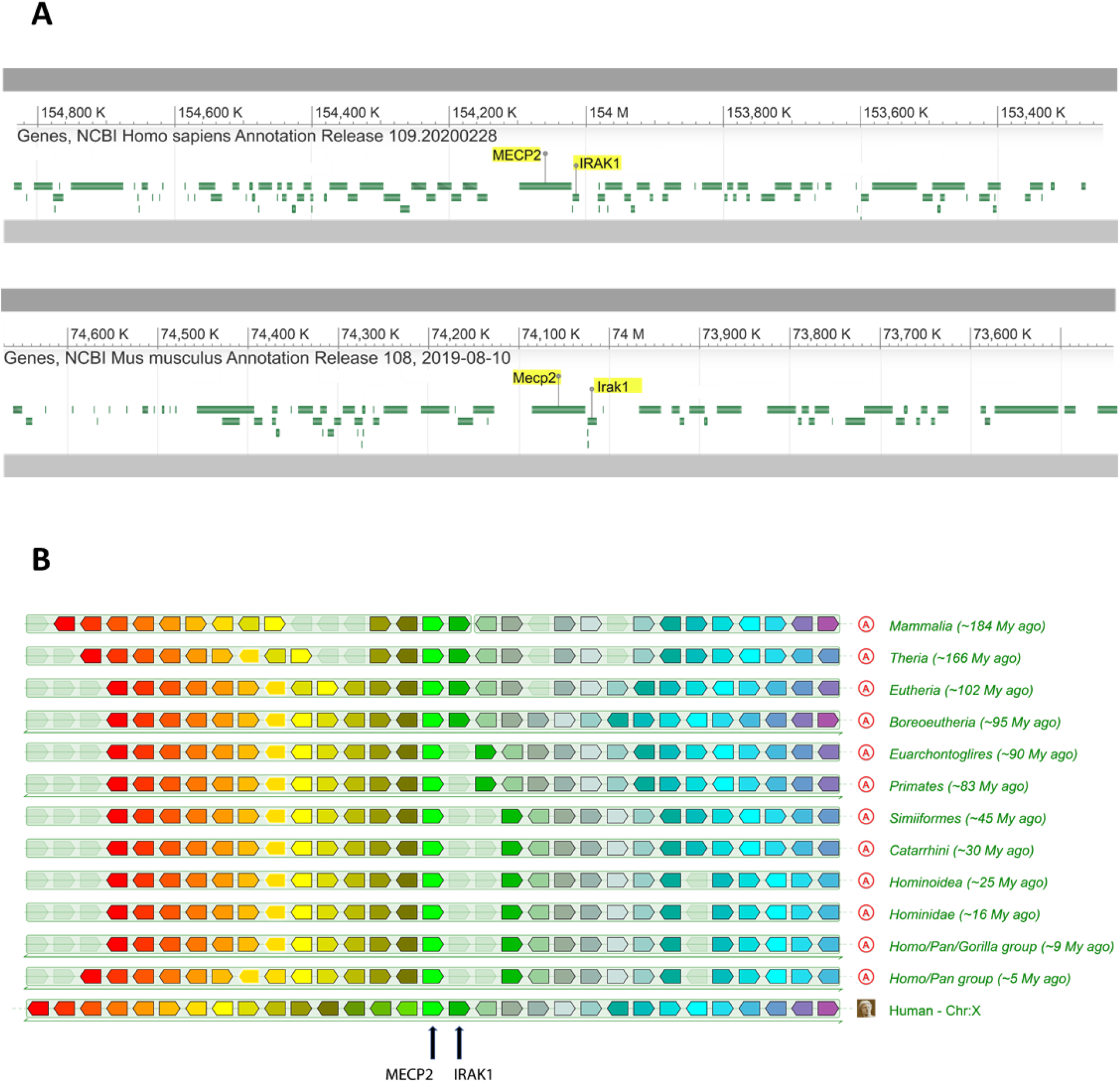
Synteny between MECP2 and IRAK1. A) NCBI Genome Data Viewer view of the genomic locations of MECP2 and IRAK1 (top panel). The gene orthologs are shown in mus musculus chromosome X (bottom panel). B) Inference of the synteny in ancestral species from Genomicus.

**Figure S4.**
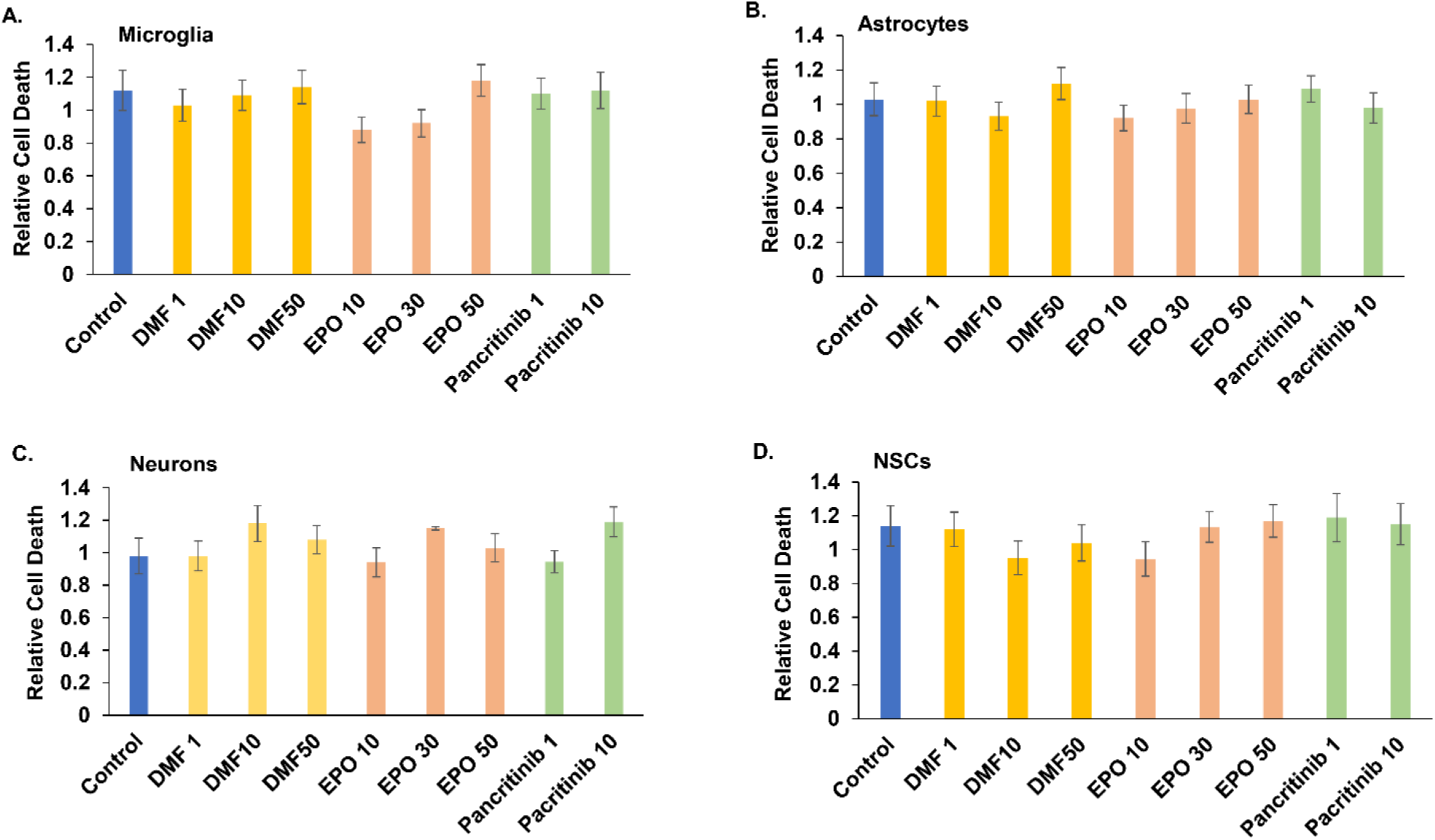
Cytotoxic effects of tested compounds. Human microglia (A), astrocytes (B), neurons (C) and neuronal stem cells (D) were treated with different concentrations of DMF (1 10 and 50 uM), EPO (10, 30 and 50 ng/ml) and pacrinitib (1 and 10 uM). Following three days of treatment, the cells were analyzed for cell death using LDH assay. The results are the means ± SD of six samples for each compound.

**Figure S5.**
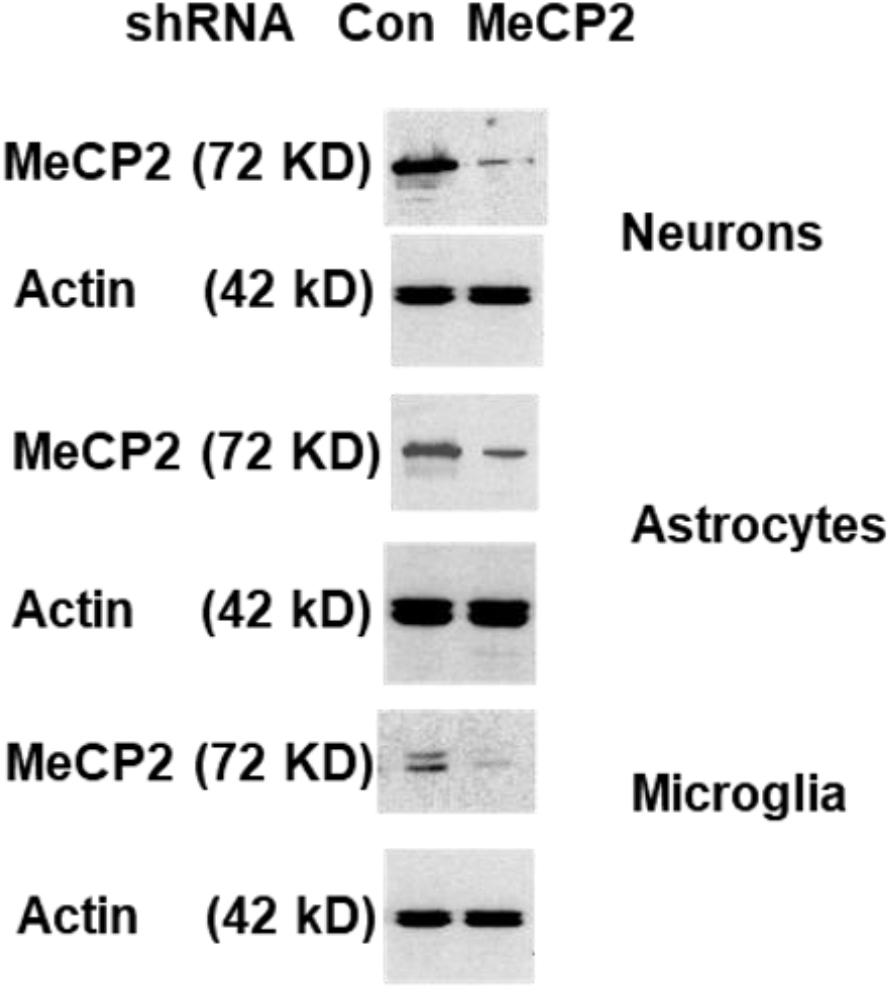
Silencing of MECP2 in neural cells. Neurons, astrocytes and microglia cells were transduced with lentivirus vectors expressing a control or MECP2 shRNAs. Following three days, the expression of MECP2 and actin were determined using Western blot analysis. The results are representative of three experiments with similar results.

**Table S1.**
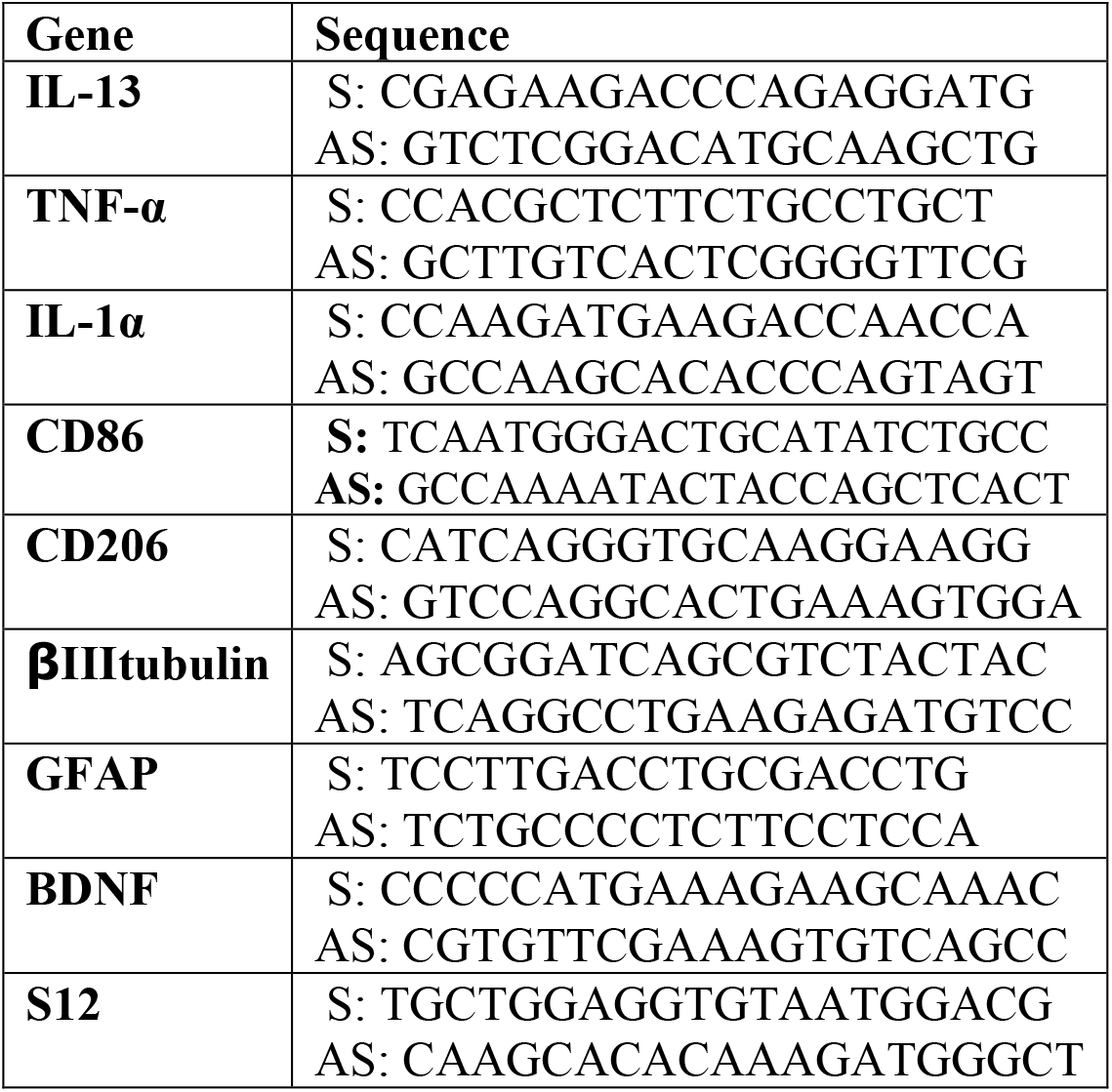
Sequences of primers used for RT-PCR.

